# A unique lower X-gate in TASK channels traps inhibitors within the vestibule

**DOI:** 10.1101/706168

**Authors:** Karin E. J. Rödström, Aytuğ K. Kiper, Wei Zhang, Susanne Rinné, Ashley C. W. Pike, Matthias Goldstein, Linus Conrad, Martina Delbeck, Michael Hahn, Heinrich Meier, Magdalena Platzk, Andrew Quigley, David Speedman, Leela Shrestha, Shubhashish M.M. Mukhopadhyay, Nicola A. Burgess-Brown, Stephen J. Tucker, Thomas Mueller, Niels Decher, Elisabeth P. Carpenter

**Affiliations:** Structural Genomics Consortium, University of Oxford, Oxford, OX3 7DQ, UK; Institute for Physiology and Pathophysiology, Vegetative Physiology, University of Marburg, Marburg, Germany; Department of Physics, University of Oxford, OX1 3QT, UK; Bayer AG, Research & Development, Pharmaceuticals, Wuppertal, Germany; CAS Key Laboratory of Pathogenic Microbiology and Immunology, Institute of Microbiology, Chinese Academy of Sciences, Beijing 100101, China; Membrane Protein Laboratory, Research Complex at Harwell, Harwell, UK

## Abstract

TASK channels are unusual members of the two-pore domain potassium (K_2P_) channel family, with unique and unexplained physiological and pharmacological characteristics. TASKs are found in neurons^1,2^, cardiomyocytes^3–5^ and vascular smooth muscle cells^6^ where they are involved in regulation of heart rate^7^, pulmonary artery tone^6,8^, sleep/wake cycles^9^ and responses to volatile anaesthetics^9–12^. K_2P_ channels regulate the resting membrane potential, providing background K^+^ currents controlled by numerous physiological stimuli^13,14^. Unlike other K_2P_ channels, TASK channels have the capacity to bind inhibitors with high affinity, exceptional selectivity and very slow compound washout rates. These characteristics make the TASK channels some of the the most easily druggable potassium channels, and indeed TASK-1 inhibitors are currently in clinical trials for obstructive sleep apnea (OSA) and atrial fibrillation (Afib)^15^ (The DOCTOS and SANDMAN Trials). Generally, potassium channels have an intramembrane vestibule with a selectivity filter above and a gate with four parallel helices below. However, K_2P_ channels studied to date all lack a lower gate. Here we present the structure of TASK-1, revealing a unique lower gate created by interaction of the two crossed C-terminal M4 transmembrane helices at the vestibule entrance, which we designate as an ‟X-gate”. This structure is formed by six residues (V^243^LRFMT^248^) that are essential for responses to volatile anaesthetics^11^, neuro-transmitters^16^ and G-protein coupled receptors^16^. Interestingly, mutations within the X-gate and surrounding regions drastically affect both open probability and activation by anaesthetics. Structures of TASK-1 with two novel, high-affinity blockers, shows both inhibitors bound below the selectivity filter, trapped in the vestibule by the X-gate, thus explaining their exceptionally low wash-out rates. Thus, the presence of the X-gate in TASK channels explains many aspects of their unusual physiological and pharmacological behaviour, which is invaluable for future development and optimization of TASK modulators for treatment of heart, lung and sleep disorders.

---

To understand the molecular basis of the unusual properties of TASK channels we solved the structure of human TWIK-related acid-sensitive potassium channel 1 (TASK-1, K_2P_3.1) by X-ray crystallography. We used a C-terminally truncated construct (Met1 to Glu259) for the structural studies, which had similar properties to the full-length channel, in particular its response to the pore-blockers A1899 and ML365, and its activation by pH (Extended Data Fig. 1). We grew crystals using the HiLiDe method^17^ and determined the structure by molecular replacement using X-ray data to 3.0 Å resolution (Methods, Extended Data Table 1, Fig. 1, Extended Data Fig. 2a-f). TASK-1 forms a domain-swapped homodimer (Fig. 1a), with a pseudo-tetrameric selectivity filter, four transmembrane helices (M1-M4) and an extracellular cap, similar to previously solved structures of other K_2P_ channels^18–22^ (Fig. 1b). The selectivity filter resembles that of other K_2P_ channels, with binding sites for four K^+^ ions. TASK-1 is inactivated by extracellular acidic pHs^23^ (Extended Data Fig. 1c) and His98, near the top of the selectivity filter, is thought to be the extracellular pH sensor^24^. Crystals were grown at pH 8.5 and the structure shows His98 forming a hydrogen bond to a water molecule behind the selectivity filter (Extended Data Fig. 2c). His98 is positioned below the sidechains of Gln77 and Gln209, which interact with each other, so that in this conformation it cannot access the extracellular space. It has been proposed that protonation of His98 could lead to reorientation of the sidechain into the extracellular space^25^, which would also require disruption of the Gln77-Gln209 interaction.

**Fig. 1.**
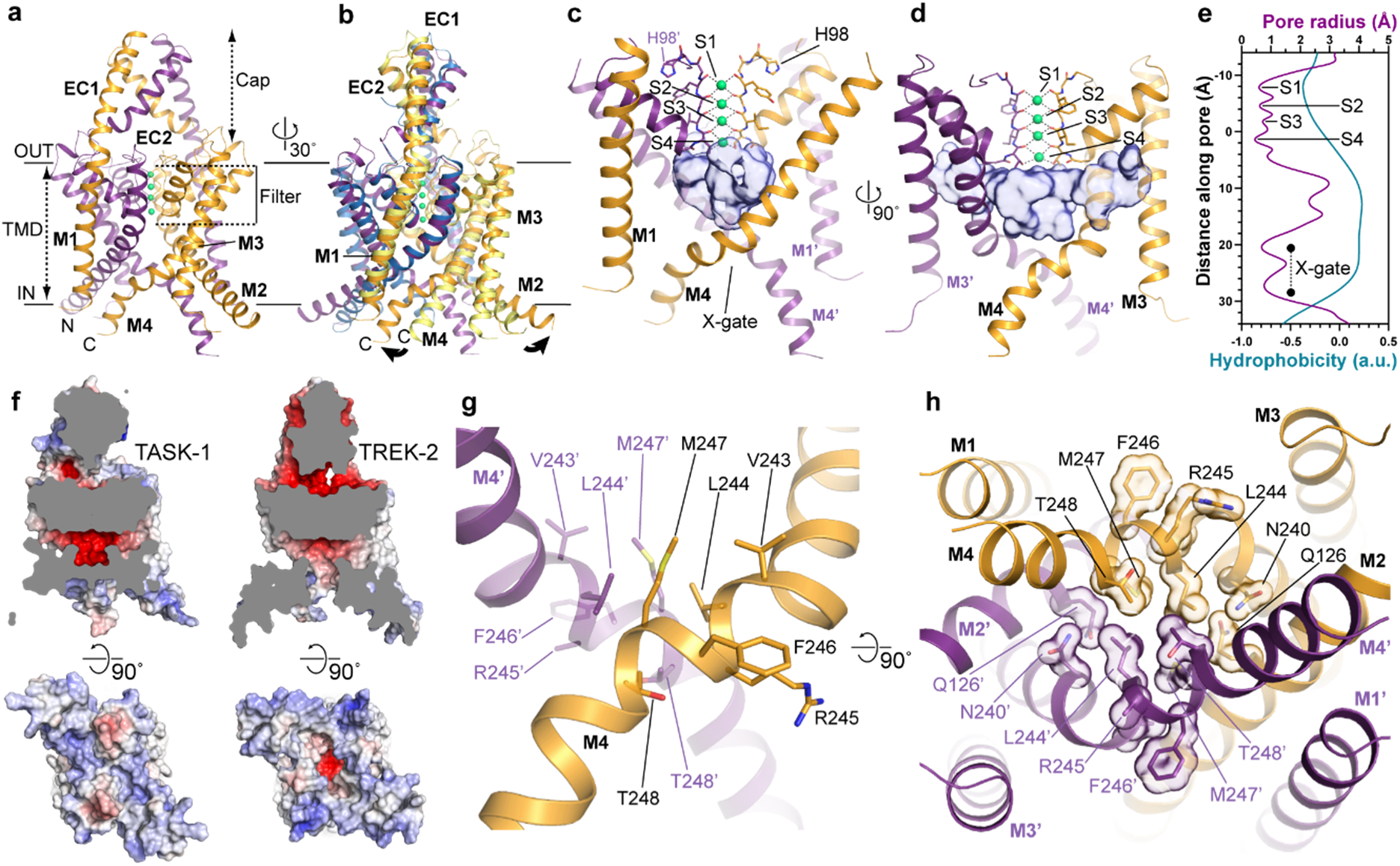
The structure of TASK-1 reveals a unique crossed helix X-gate. **a**, Cartoon of the TASK-1 structure, with chain A (gold), B (purple) and potassium ions (green), viewed from the membrane plane. **b**, Superposition of the TASK-1 and TREK-2 (yellow and blue) “down” state (PDB: 4XDJ) structures. The TASK-1 structure with the X-gate enclosing the vestibule and fenestrations (depicted with semi-transparent surfaces), **c**, viewed from the membrane plane and **d**, rotated 90°. **e**, Radius and hydrophobicity of the TASK-1 pore (CHAP^49^). **f**, Electrostatic surfaces of TASK-1 and TREK-2 (PDB: 4XDJ) shown as a cross section perpendicular to membrane and seen from below. The X-gate viewed from **g**, the plane of the membrane and **h**, below the membrane.

One unusual feature of the TASK and THIK channels is the absence of the intermolecular disulphide bond at the apex of the external cap domain (Extended Data Fig. 3b). However, we found that in TASK-1 the domain-swapped cap structure is preserved without this disulphide bond (Extended Data Fig. 2e-h), as previously predicted^26^. Therefore, this feature does not explain the differences in TASK-1 properties. The relative positions of the M2-M4 helices determine whether stretch-activated K_2P_ channels are in a more active “up” state with raised helices and closed fenestrations, or a lower activity “down” state with open fenestrations^20,21^. Overall the TASK-1 structure adopts a “down”–like state, with some adjustment of the M2 position and open fenestrations connecting the small central vestibule to the lipid bilayer (Fig. 1b-d).

Surprisingly, we found that at the distal end of M4 there was an unexpected conformational rearrangement, causing the M4 helices to lie across the entrance of the vestibule, forming a closed gate which we have designated as an “X-gate” (Fig. 1c-h). This conformational change is caused by bends in M4 at residues Leu241 and Asn250, on either side of a six residue sequence (V^243^LRFMT^248^) which forms the X-gate. This leaves a gap of 2-3 Å between the M4 helices, in contrast to TREK-2 which has a gap of at least 7.5 Å (Fig. 1e,f). The X-gate adopts an unusual extended two-turn helical structure, lacking the hydrogen bonding patterns seen in α or 3_10_ helices (Extended Data Fig. 2i). The interface is lined by the hydrophobic sidechains of Leu244, Met247 and the methyl group of Thr248 (Fig. 1g,h), which block the ion conduction pathway. Interestingly, this region is essential for TASK-1 responses to volatile anaesthetics^9–12^, neurotransmitters^16^, drugs^27^ and G-protein coupled receptors^16^ (potentially through diacylglycerol^28^ or PIP_2_ interactions^29)^. Sequence alignments suggest TASK-3 and TASK-5 may also form similar structures (Extended Data Fig. 3). The presence of the X-gate in the TASK subfamily is an exception to the paradigm that K_2P_ channels lack a lower gate^30^. Surprisingly, the structure of the dimeric X-gate is unrelated to the ‘helix bundle crossing’ gates found in tetrameric K^+^ channels, which have four parallel helices forming a funnel shape.

To determine whether the X-gate explains the unusual properties of TASK channels, we performed alanine scanning mutagenesis across the distal M4 helix (Thr230-Thr248), and assessed channel activity and surface expression of the mutants. We predicted that disruption of the X-gate would increase the very low TASK-1 basal channel activity^7,30,31^. Strikingly, mutation of any residue within the X-gate (except Val243), led to increased currents (Fig. 2a), which were not caused by increased surface expression, as we observed similar or reduced surface expression (Extended Data Fig. 4a). The centre of the X-gate is formed by Leu244, which is packed between residues on M2, M4 and M4’ (Fig. 1h,2b) and the Val244A mutation resulted in the largest current increase (Fig. 2a). Inside out single channel recordings of Leu244Ala confirmed a direct effect on channel gating with a 6-fold increase in channel open probability from 0.7 to 4.0% (Fig. 2c,d) and increased channel open times (Fig. 2e). Mutation of the equivalent residue in TASK-3 (Leu244Ala) also resulted in increased channel activity (Fig. 2f). The X-gate is also important for activation by volatile anaesthetics such as halothane and sevoflurane^9–12^, and was originally called the “halothane response element”. Here we confirm that sevoflurane, a modern and widely used anaesthetic, also activates TASK-1 and that mutations in the X-gate reduces this activation (Fig. 2g,h), indicating a role for the X-gate in responses to volatile anaesthetics.

**Fig. 2.**
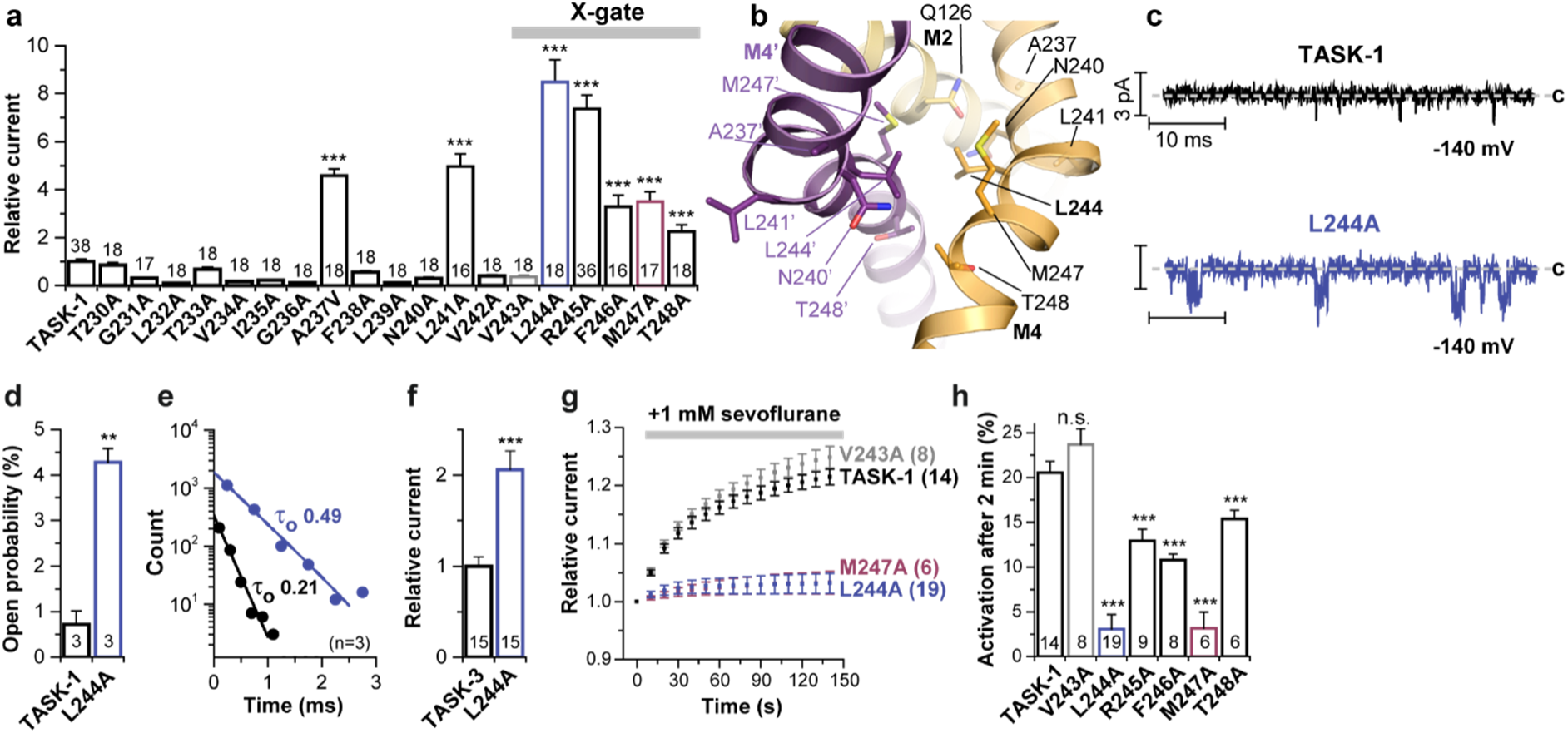
Mutation of X-gate residues results in an increase in ion conduction and loss of anaesthetic activation. **a**, Relative current amplitudes of wild-type and mutated TASK-1. **b**, The X-gate viewed from the membrane plane, rotated 90° relative to Fig. 1g showing Leu244 and surrounding residues. **c**, Representative inside-out single channel recordings of TASK-1 wild-type and Leu244Ala. Analysis of **d**, the open probability and **e**, the mean open times. **f**, Relative current amplitudes of wild-type TASK-3 and Leu244Ala. **g**, Application of 1 mM sevoflurane to wild-type and mutated TASK-1 and **h**, analysis 2 minutes after application. All two-electrode voltage-clamp recordings and single channel measurements in Fig. 2–4 were performed in *X. laevis* oocytes. The number of experiments performed are illustrated in the respective graphs. Panels **a,d,f,g,h** data are presented as mean ± s.e.m..

Next we investigated residues surrounding the X-gate, which may be important for its formation and stability. The X-gate is preceded by a hinge point in M4 around Leu241-Val242, where the helix axis is bent by ~40° (Fig. 3a). After the bend the X-gate itself adopts an expanded helical conformation (Extended Data Fig. 2i). The side chain of Leu241 inserts into a hydrophobic pocket, pulling it away from the helix axis and stabilizing the bend in M4. This exposes the Leu241 carbonyl so that it can hydrogen bond with the Arg245 sidechain (Fig. 3a). Mutation of either Leu241 or Arg245 to Ala increased channel activity by 5-7 fold (Fig. 2a), confirming the importance of these interactions in X-gate stability. Furthermore, a hydrogen bond between the carbonyl oxygen of Ala237 on M4 and Asn133 on M2 also disrupts the M4 α helix, stabilizing the bend, and mutation of Asn133 increased channel activity (Fig. 3b).

**Fig. 3.**
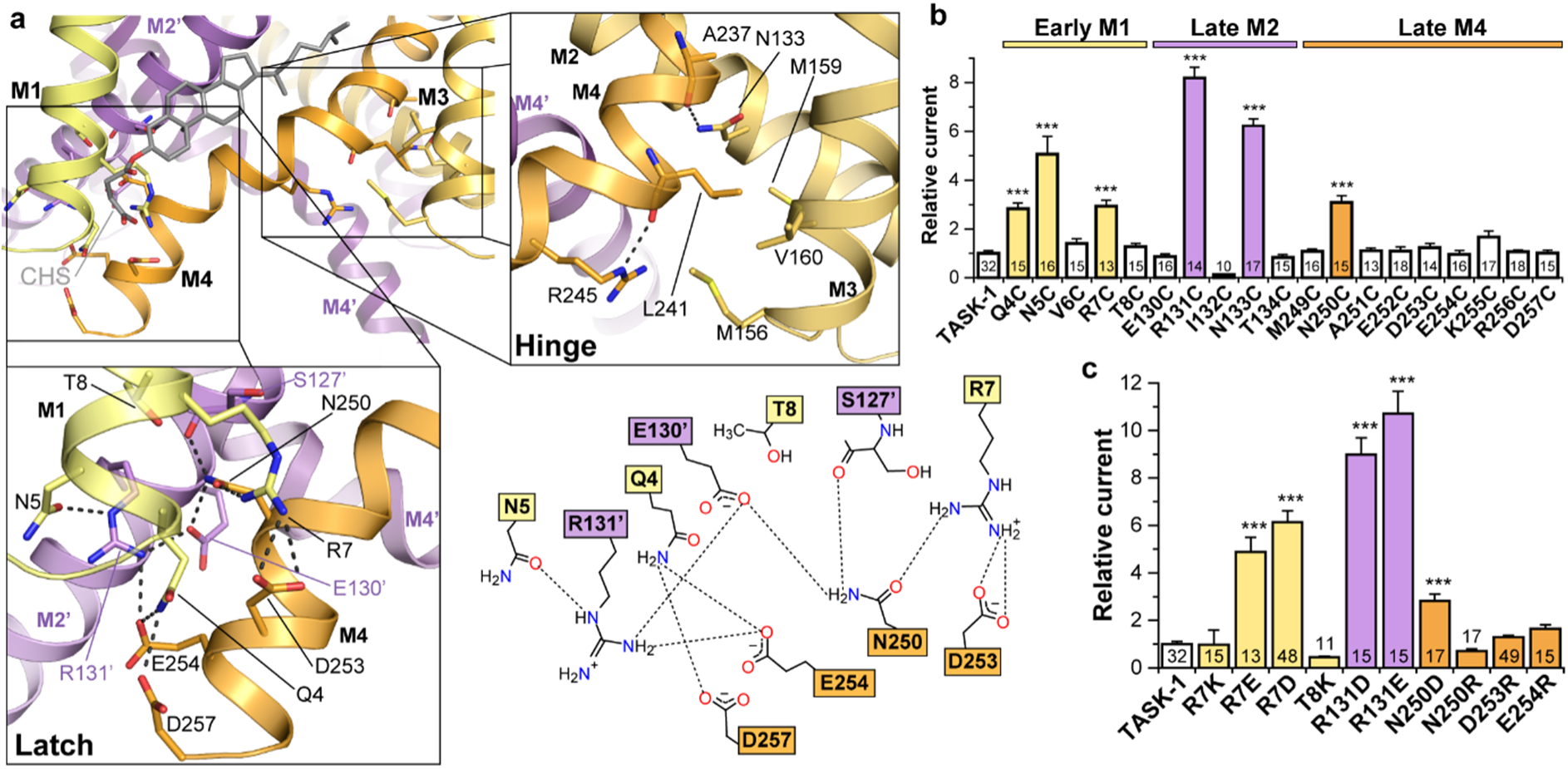
The X-gate environment includes a hinge and a latch on either side of the vestibule which are required for X-gate integrity. **a**, The X-gate in TASK-1, with the cholesteryl hemisuccinate (CHS) binding site. The hinge is shown on the right and the latch region below. **b**, Relative current amplitudes of wild-type and mutated TASK-1. **c**, Relative current amplitudes for mutated residues in the latch. **b, c**, Data are presented as mean ± s.e.m..

Following the X-gate, a second bend in M4 at residue Asn250 allows the distal end of M4 to adopt an α-helical structure which forms extensive interactions with the opposite side of the vestibule, a region we have named the “latch” (Fig. 3a). This consists of an extensive network of salt bridges and hydrogen bonds linking M4 to M1 and M2’ (Fig. 3a). Mutation of several residues across the proximal M1, distal M2’ and distal M4 regions gave increased channel activity, consistent with the latch region playing an important role in stabilizing the X-gate (Fig. 3b). In particular, Arg7 and Arg131’ appear to be important for the integrity of the latch, since mutations to acidic residues causes a 5 – 10 fold increase in currents (Fig. 3c). The reduced levels of surface expression for these mutants indicates their increased currents are likely due to direct effects on channel gating, not increased surface expression (Extended data Fig. 4b).

Lipids are also important regulators of K_2P_ channels and the structures reveal binding sites for lipids, including the cholesterol analogue cholesteryl hemisuccinate (CHS) (Extended Data Fig. 2d). CHS (added during TASK-1 purification) extends along the X-gate, interacting with M1, M4 and M2’ (Fig. 3a). Phe246 in the X-gate lies next to the CHS molecule (Fig. 3a), and the Phe246Ala mutation produces a 4-fold increase in current (Fig. 2a). Therefore, cholesterol may also help to stabilize the X-gate in a closed conformation.

Unlike many other K_2P_ channels that exhibit very poor pharmacology, several high affinity and highly selective inhibitors exist for TASK-1 and TASK-3 channels. Furthermore, these blockers exhibit exceptionally low wash-out rates after removal of the inhibitor. Together, these characteristics make TASK inhibitors interesting pharmacological tools to investigate the physiological and pharmacological function of TASK channels. Therefore, we performed an ultra-high throughput screen (uHTS) on TASK-1, which identified two highly potent and selective inhibitors of TASK-1, BAY 1000493 and BAY 2341237 (Fig. 4a,b), with EC_50_ values of 9.5 and 7.6 nM. Crystal structures of complexes of TASK-1 with these inhibitors showed that they both bind within the inner vestibule, directly below the selectivity filter (Fig 4c-g, Extended Data Fig. 5). In both cases, a single molecule binds across the 2-fold axis of the dimer, with one molecule bound per dimer. The electron density for BAY 1000493 shows a superposition of two copies of the inhibitor, with 50% occupancy for each orientation (Extended Data Fig. 5a,d), as expected. This arrangement was confirmed using anomalous difference maps with data collected at the Br absorption edge (Extended Data Fig. 5a). In contrast, BAY 2341237 only binds in one orientation, with 100% occupancy, due to small differences in the two chains of the dimer (Fig. 4d,f,g, Extended Data Fig. 5b,e).

**Fig. 4.**
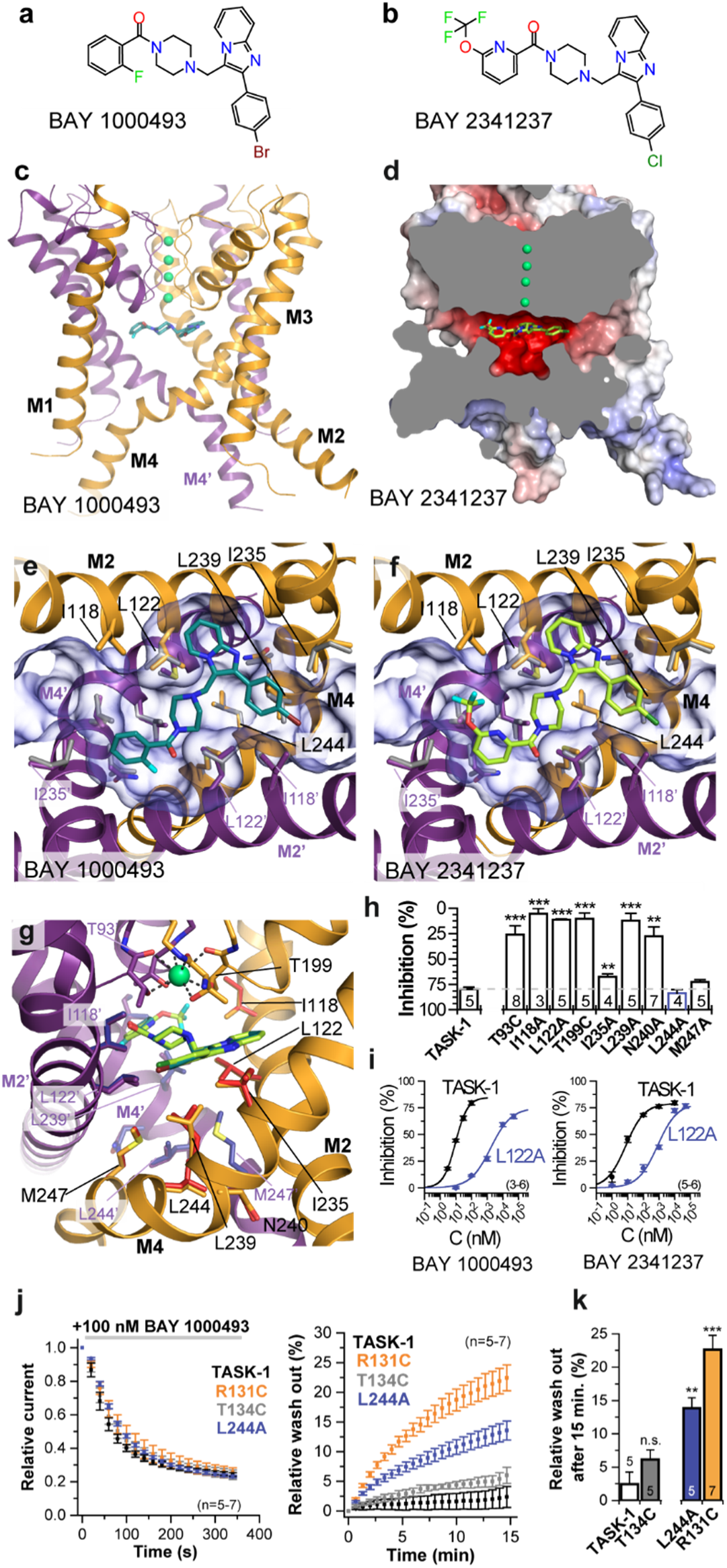
Structures of complexes with two highly potent inhibitors show binding in the vestibule, trapped by the X-gate. Chemical structures of **a**, BAY 1000493 ((4-{[2-(4-bromophenyl)imidazo[1,2-a]pyridin-3-yl]methyl}vpiperazin-1-yl)(2-fluorophenyl)methanone) and **b**, BAY 2341237 (4-{[2-(4-chlorophenyl)imidazo[1,2-a]pyridin-3-yl]methyl} piperazin-1-yl)[6-(trifluoromethoxy) pyridin-2-yl]methanone. **c**, Structure of the TASK-1/BAY 1000493 (teal) complex (M2’ excluded). For simplicity, only one orientation of BAY 1000493 is shown, although it binds to the dimer equally well in either orientation (see Extended Data Fig. 6a). **d**, Cross section of an electrostatic surface of the TASK-1/BAY 2341237 (green) complex. **e**, The BAY 1000493 and **f**, BAY 2341237 binding sites below the selectivity filter, looking towards the X-gate and **g**, superposition of the two complexes seen from the membrane plane with sidechains from the BAY 2341237 complex in red and blue. **h**, Percentage of inhibition of wild-type and mutated TASK-1 by 100 nM BAY 1000493. **i**, Dose-response curves for BAY 1000493 and BAY 2341237 for wild-type TASK-1 and Leu122Ala. **j**, Analysis of relative current amplitudes over time while washing BAY 1000493 in and out. **k**, Analysis of the relative wash-out. Data in **h,i,j,k**, are presented as mean ± s.e.m..

Both inhibitors bind to the closed conformation of TASK-1, below the selectivity filter, without perturbing the X-gate structure. They fit precisely to the surface at the top of the vestibule, with Leu122 projecting into the vestibule below the compounds, holding them in place (Fig. 4e-g). Consistent with these observations, we found that mutation of residues lining the inhibitor binding site (Thr93Cys, Ile118Ala, Leu122Ala, Thr199Cys and Leu239Ala) drastically reduced inhibition by BAY 1000493 (Fig. 4h). Leu122 is important for binding of other TASK-1 pore blockers^27,32^ and it occupies a key position below the binding site for our inhibitors (Fig. 4g). When we recorded full dose-response curves with TASK-1 and a Leu122Ala mutant, we found a >100-fold decrease in EC_50_ values for both inhibitors with the mutation (Fig 4i). In addition, we found that BAY 1000493 is highly selective for TASK-1 and TASK-3, when compared to a wide range of other ion channels (Extended Data Fig. 6a,b, Extended Data Table 2).

Our structures show that both compounds bind at the top of the vestibule, trapped but not in direct contact with the X-gate. Leu244 is central to the X-gate structure, but is not part of the inhibitor binding site and a Leu244Ala mutation did not alter inhibition by BAY 1000493 (Fig. 4h). However, TASK channels have another critical feature, their ability to retain bound compounds for long periods during washout experiments. BAY 1000493 shows very slow washout rates from TASK-1 (Fig. 4j, Extended Data Fig. 6c), unlike a larger TASK inhibitor, A1899, which has an IC_50_ value of 35 nM and washes off significantly faster^27^ (Extended Data Fig. 6d-f). When the Leu244Ala mutation is introduced at the X-gate or the Arg131Asp mutation at the latch, we observed a significant increases in the BAY 1000493 washout rate (Fig. 4j,k, Extended Data Fig. 6c). Conversely, a mutation near the latch (Thr134Cys), that has relatively little effect on X-gate function (Fig. 3b), also has less effect on washout rates (Fig., 4j, Extended Data Fig. 6c). This suggests that disruption of the X-gate or the latch may stabilise the open state of the channel, thereby allowing BAY 1000493 to dissociate more easily from this site.

The X-gate must be capable of opening wide enough to allow inhibitors to enter, as well as hydrated K^+^ ions. Therefore, it is likely that X-gate opening requires a significant conformational change, not simply small adjustments in sidechain positions or helix orientation. We speculate that this could involve opening of the latch, relaxation of the X-gate extended helix into an α-helical conformation and straightening of M4 (Extended Data Fig. 7), giving a conformation similar to other K_2P_ structures^18–22^. This would also allow the M1-M4 helices to adopt a more “up”-like conformation, without fenestrations, consistent with the proposed higher activity state of other K_2P_s (Extended Data Fig. 7).

Loss of function (LoF) missense variants in TASK channels are associated with two genetic diseases. Mutations in TASK-1 cause autosomal dominant primary pulmonary hypertension Type 4^33^ (PPH4: OMIM: 615344), a serious condition that is often fatal in mid-life^33^, whilst mutations in TASK-3 are linked to the developmental disorder Birk-Barel syndrome^34^. These mutations are distributed throughout the structure (see Extended Data Fig. 8a-g, Supplementary Discussion). However, one PPH4-associated TASK-1 mutation, Thr8Lys on M1, is located within the X-gate latch and causes ~2-fold decreased currents (Fig. 3a, Extended Data Fig 8c). Furthermore, two Birk-Barel syndrome associated mutations in TASK-3 (Gly236Arg^34^ and Ala237Asp^35^) preceded the hinge on M4. Gly236Arg places a large positively charged residue within the vestibule, near the ion conduction pathway and the X-gate, whereas Ala237Asp places a charged residue adjacent to the hinge (Extended Data Fig. 8h). These mutations emphasize the significance of these regions in TASK channel function.

In conclusion, we have shown that the TASKs channels are highly unusual members of the K_2P_ channel family in that they have the ability to form a cytoplasmic X-gate, which may contribute to the low open probability of TASK-1. We have also shown that two nanomolar affinity inhibitors bind in the closed vestibule, trapped by the X-gate. Their precise fit to the top surface of the vestibule, combined with the trapping effect of the X-gate may explain the exceptionally high affinity and selectivity observed with these pore blockers. In addition, the presence of the X-gate may also explain the very low wash-out rates of these inhibitors. Compound trapping within the inner cavity has been proposed previously for many ion channels^36–39^. TASK-1 inhibitor binding requires opening of the X-gate, allowing compounds to enter and bind to the high affinity site below the selectivity filter. Since the X-gate is closed most of the time and there is little space for the compound to dissociate, they will remain bound, trapped by the X-gate, resulting in an increased apparent affinity and reduced washout rate. However, whether the open state of TASK-1 involves any change in the conformation of the compound binding site, or whether these compounds can stabilize the X-gate in its closed conformation, remains to be seen.

The X-gate is clearly important for the control of TASK channel activity and may contribute to their regulation by neurotransmitters^16^, intracellular signalling pathways^16,28,29^, volatile anaesthetics^9–12^ and small molecules^27^. However, given the polymodal nature of K_2P_ channel regulation and the known role of the selectivity filter in K_2P_ channel gating, the X-gate is unlikely to be the only route for TASK channel regulation. For example, their physiologically relevant regulation by extracellular pH involves His98, which is extracellular, located next to the selectivity filter. Further studies are needed to determine if these two gating mechanisms are coupled.

This study also provides insight into the mechanism of TASK-1 inhibitors that are already in clinical trials as potential treatments for obstructive sleep apnea and atrial fibrillation. The unique environment of the inhibitor binding site within the vestibule, created by the X-gate, has significant implications for interpretation of pharmacokinetic properties of TASK inhibitors. In particular, questions of bioavailability, metabolism and half-life should be interpreted in the light of the discovery of the X-gate. Once a blocker becomes sequestered within this vestibule, it is likely to remain bound and ‘active’ for much longer than anticipated by studying its pharmacokinetic properties. Such factors should be carefully evaluated during drug development and trials. Overall, this work provides critical understanding of the gating mechanism in TASK channels and the action of clinically-relevant inhibitors, processes that are of uppermost importance for their use as targets for efficient and safe treatment of disease.

## Methods

### Cloning and expression of TASK-1^Met1-Glu259^ for structural studies

The human *KCNK3* gene (Genbank ID 4504849), which encoded the TASK-1 protein, was obtained from Origene. The TASK-1 construct, residues Met1 to Glu259, was subcloned in the SGC vector pFB-CT10HF-LIC, with a C terminal TEV site followed by decahistidine and FLAG tags. Baculovirus was generated with the Bac-to-bac system and passaged twice before it was used for infecting Sf9 cells at a density of 2 × 10^6^ cells/mL and a ratio of 5 mL virus per litre cells. Infected cells were harvested after 72 h by centrifugation, frozen in liquid nitrogen and stored at −80°C.

### Purification of TASK-1

Cells were broken with an EmulsiFlex-C3 or C5 high-pressure homogenizer (Avestin Inc.) in 40 ml breaking buffer (50 mM HEPES pH 7.5, 200 mM KCl, 5% v/v glycerol) per initial litre culture volume. Protein was solubilized by addition of 1% w/v n-decyl-β-D-maltopyranoside (DM) (Anatrace) and 0.1% w/v cholesteryl hemisuccinate (CHS) Tris salt (Sigma-Aldrich) and rotated at 4°C for 1 h. Cell debris was pelleted at 35,000 g, 1 h, 4°C, 5 mM imidazole pH 8.0 was added, and protein was incubated with Talon (Clontech), 0.5 ml resin per litre cell culture for 1 h at 4°C. Resin was collected and washed with 30 CV wash buffer (50 mM HEPES pH 7.5, 200 mM KCl, 5% v/v glycerol, 20 mM imidazole pH 8.0, 0.24% w/v DM, 0.024% w/v CHS), and eluted with elution buffer (50 mM HEPES pH 7.5, 200 mM KCl, 5% v/v glycerol, 250 mM imidazole pH 8.0, 0.24% w/v DM, 0.024% w/v CHS). The protein was exchanged into desalting buffer (50 mM HEPES pH 7.5, 200 mM KCl, 5% v/v glycerol, 0.24% w/v DM, 0.024% w/v CHS) by passing over PD10 columns and purification tags were cleaved by addition of a 1:3 w:w ratio of 6x His-tagged TEV protease and deglycosylated with a 1:10 w:w ratio of 6x His-tagged PNGaseF, for 12-16 h at 4°C. Enzymes were removed by addition of 0.1 ml Talon resin per litre initial cell culture and 7.5 mM imidazole pH 8.0. The protein was concentrated to 500 µl and subjected to size exclusion chromatography on a Superose 6 Increase 10/300 GL column (GE Healthcare) in gel filtration buffer (20 mM HEPES pH 7.5, 200 mM KCl, 0.12% w/v DM, 0.012% w/v CHS). Fractions containing the highest concentration of TASK-1 were pooled and concentrated to 12-30 mg/ml ml with a 50 kDa molecular weight cut off spin concentrator (GE Healthcare).

### Crystallisation, X-ray data collection and data processing

Crystallization trials were performed in 96-well sitting drop vapour diffusion plates using a Mosquito crystallization robot (TTP Labtech) at 8 - 12 mg/ml with 150 nl drops and 2:1, 1:1 and 1:2 protein:reservoir ratios. Initial crystals were obtained at 20°C in an in-house version of MemGold HT-96, condition D7 (0.1 M TRIS pH 8.5, 0.1 mM KCl and 39% v/v polyethylene glycol (PEG) 400). Crystals were further optimized by addition of 3% w/v sucrose to the precipitant solution and by using the HiLiDe method^17^. Briefly, 1,2-dioleoyl-sn-glycero-3-phosphocholine (DOPC) (Avanti Polar Lipids, Inc.) in chloroform was dried down in a round-bottomed glass vial, 1.5 µg lipid per µl protein. Protein at 5.7-6.2 mg/ml was added along with 15 µg/µl DM and 1.5 µg/µl CHS, incubated slowly shaking at 4°C for 16-24 h and centrifuged at 15,000 g, 2 h, 4°C. Crystallization was carried out in 24-well hanging drop plates with 2 µl drops and a 2:1 protein:reservoir ratio at 20°C. Crystals grew over 1-8 weeks in 0.1 M TRIS pH 8.5, 0.05 M KCl, 32% PEG 400 and 3% w/v sucrose and were harvested at 6°C directly from the drop and vitrified.

Data collection was carried out at Diamond Light Source Ltd. (DLS) using the microfocus beamline I24. The final dataset was assembled from a series of 40° wedges collected from two crystals, using a 9×6 µm microbeam.

Crystals of complexes of TASK-1 with inhibitors were also grown using the HiLiDe method. Each inhibitor, dissolved in 100% DMSO at 130 mM, was added to the HiLiDe setup to a final concentration of 1.3 mM. Crystals grew in 0.1 M TRIS pH 8.5, 0.05 M KCl, and 3% w/v sucrose with 24% PEG 400 for BAY 1000493 and 31% PEG 400 for BAY 2341237 over 1 - 12 weeks and were harvested at 6°C. Data were collected at the DLS on beamline I24 at a wavelength of 0.9686 Å, with 0.1 s exposures, a beam size of 20×20 µm and 0.2° oscillations. A full dataset for the BAY 1000493 complex was assembled from several 40° wedges from one crystal whereas the BAY 2341237 dataset was collected as a single wedge, both with a 20×20 µm beam. Anomalous data for BAY 1000493 were collected at the bromine edge, (λ = 0.9116 Å) in four 360° passes.

### Model building and refinement for structures

Data were processed as individual wedges in XDS^40^ and scaled together in XSCALE^40^. The data were merged in AIMLESS^41,42^ (Extended Data Table 1). Due to the anisotropy of the data, an anisotropic cut-off was applied using the STARANISO server^43^. The structure was solved using molecular replacement with Phaser, using a truncated model of TREK-2^21^ (PDB ID: 4BW5) with separate search models consisting of the TM domain and the cap. Two copies of the TASK-1 dimer were located in the asymmetric unit. An initial TASK-1 model was built using a density modified prime-and-switch map calculated using *phenix.autobuild*^44^ as a guide and was improved by several rounds of manual model building and refinement in COOT^45^. Refinement was carried out in BUSTER version 2.10.3^46^, using all data to 3.0 Å in the highest resolved direction, with NCS restraints and one TLS group per chain. Four potassium ions per homodimer were modelled and distances from the oxygen atoms in the filter were restrained to 2.8 Å. The final model for the native structure comprised residues Met1 to Asp257 in chains A and C, and Met1 to Leu261 in chains B and D. In the AB dimer, residues 149 to 151 are disordered and were not modelled. The selectivity filter of TASK-1 consists of residues T^93^IGY^96^ and T^199^IGF^202^ from each subunit, and they adopt the expected conformation seen in other K_2P_ structures^18–22^. The carbonyl oxygens of these residues, as well as the sidechain oxygens of Thr93 and Thr199, coordinate four K^+^ ions in sites S1-S4 (Fig. 1c,d, Extended Data Fig. 2a-d). However, there is weaker density for the S2 K^+^ ion and it refines to have a higher B factor in all structures, compared to the three other K^+^ ions. Omitting this K^+^ ion altogether resulted in a positive Fo-Fc peak, so it was included in the model, with 100% occupancy and a higher B factor.

The structure contains a well-defined CHS molecule packed against the X-gate. It lies in an extensive, largely hydrophobic groove formed by residues Arg3, Arg7, Pro119, Val123, Val243, Phe246 and Met249. The two arginine residues are adjacent to the hemisuccinate moiety whereas the hydrophobic residues are adjacent to the cholesterol unit.

Data for the ligand complexes were processed in the same way as the native complex (Extended Data Table 1) to 2.9 Å and 3.1 Å for BAY 1000493 and BAY 2341237, respectively. After one round of refinement in BUSTER version 2.10.3^46^, positive difference density was apparent in the vestibule. For BAY 1000493, an anomalous difference map was calculated in PHENIX^44^, revealing two Br peaks per dimer. Since BAY 1000493 binds across the dimer axis below the selectivity filter, only one copy of the inhibitor is bound per TASK-1 dimer. In the case of BAY 1000493, the dimer is symmetrical, and the inhibitor binds in either orientation with a 50% probability, so the density in the pore is the average of the two possible orientations throughtout the crystal. However, each dimer has one molecule of BAY 1000493 bound and it only binds in one position, with the same interactions regardless of the orientation. In contrast, The Fo-Fc difference density map calculated with BAY 2341237 omitted from the structure, revealed a single conformation for the compound, which was modelled with 100% occupancy. This is the result of BAY 2341237 inducing some small degree of asymmetry into the dimer, resulting in the dimer/inhibitor complex packing in only one orientation in the crystal. The final models were refined against the anisotropically truncated STARANISO^43^ data.

### Two-electrode voltage-clamp measurements

*X. laevis* oocytes were obtained and the TEVC measurements were recorded as described previously^27^. Briefly, collected oocytes were stored at 18 °C in ND96 solution (in mM: NaCl 96, KCl 2, CaCl_2_ 1.8, MgCl_2_ 1, HEPES 5 and pH 7.5) supplemented with 50 mg/l gentamycin, 274 mg/l sodium pyruvate and 88 mg/l theophylline. Oocytes were injected with 5 ng of TASK-1 cRNA and incubated for 48 h at 18 °C. ND96 was used as recording solution. Oocytes were held at −80 mV and voltage was ramped from −120 to +45 mV within 3.5 s, using a sweep time interval of 4 s. Block was analysed with voltage steps from a holding potential of −80 mV. A first test pulse to 0 mV of 1 s duration was followed by a repolarising step to −80 mV for 1 s, directly followed by another 1 s test pulse to +40 mV. The sweep time interval was 10 s. All inhibitors were dissolved in DMSO, aliquoted, stored at −20 °C or room temperature and added to the external solution (ND96) just before the recordings. The EC_50_ was determined from Hill plots using four concentrations for each construct. Final DMSO concentration of 0.1 % was not exceed.

For measurements of the TASK-1^Met1-Glu259^ currents shown in the Extended Data Fig. 1 oocytes were injected with 1 ng of Task-1 RNA and two electrode voltage clamp experiments performed with a Warner oocyte-clamp (OC-725-C). Data were digitalised with an Digidata1400A interface (Molecular Devices). Voltage-ramp protocols were used to measure IV-responses. The ramps lasted for 0.5 s with ranges of −120 mV to 50 mV. In each experimental condition 10 ramps were recorded with the last 5 sweeps of such a series averaged and used for analysis. Voltage-clamp recordings were performed in ND96 solution adjusted to the desired pH at room temperature with NaOH. For experiments at pH 5.5, HEPES was exchanged for MES.

### Chemiluminescence assay in *X. laevis* oocytes

Surface expression of TASK-1 channel constructs was studied as previously described^47^. Briefly, *X. laevis* oocytes were injected with cRNA of HA-tagged channels. After 48h, oocytes were incubated in ND96 plus 1 % BSA on ice for 30 min to reduce unspecific antibody binding. Subsequently, the oocytes were incubated with primary antibody (rat anti-HA (Roche), 1:100) for 1 h, and after extensive washing they were incubated with secondary antibody (goat anti-rat-IgG, HRP-coupled (Dianova), 1:500) for 1 h. After oocytes were individually placed into a vial with 20 µl of luminescence substrate (SuperSignal Femto (Thermo Scientific)), light emission was detected with a GloMax luminometer (Promega).

### Single channel measurements

Inside-out single channel patch clamp recordings of *Xenopus* oocytes were performed similar as previously described^31^. Briefly, all experiments were conducted at room temperature with a bath solution containing in mM: KCl 140, HEPES 5, EGTA 1 and pH 7.4. Single channel current recordings were executed with an Axopatch 200B amplifier (Axon Instruments), a Digidata 1550B A/D converter (Axon Instruments) and pClamp10 software (Axon Instruments). The sampling rate was 15 kHz with the analog filter set to 2 kHz. Voltage pulses were applied from 0 mV (holding potential) to +140 mV and/or −140 mV for 1 s with an interpulse interval of 8 s. Data were analysed with ClampFit10 and Origin 2016 (OriginLab Corporation).

### Quantification and statistical analysis

Quantification and the statistical analysis of the data were executed as previously described^48^. Briefly, normality and variance of every dataset was tested. A paired or unpaired Student’s t-test was used to probe the significance but for not normally distributed data, a non-parametric Mann-Whitney U-test and a Wilcoxon signed-rank test for paired analyses was used, respectively. In case of different variances, significance was probed with Welch’s t-test and for not normally distributed data with Mood’s median test. All data are presented as mean ± s.e.m. and all experiments were repeated from N = 2 - 5 different batches (biological replicates). The number of experiments (n) as technical replicates are indicated in the subpanels of the Figures. Significances are indicated with *, P < 0.05; **, P < 0.01; ***, P < 0.001 in the Figures.

### Synthesis of TASK-1 inhibitors BAY 1000493 and BAY 2341237

The synthesis of BAY 1000493 and BAY 2341237 are described in patent number WO2017097792A1 and in the supplementary methods file.

## Acknowledgements

K.E.J.R. and D.S. are supported by B.B.S.R.C grant no BB/N009274/1. K.E.J.R., A.C.W.P., D.S., S.M.M.M., N.B.-B., E.P.C. are members of the SGC, a registered charity (number 1097737) that receives funds from AbbVie, Bayer Pharma AG, Boehringer Ingelheim, the Canada Foundation for Innovation, Genome Canada, Janssen, Merck KGaA, Merck & Co., Novartis, the Ontario Ministry of Economic Development and Innovation, Pfizer, São Paulo Research Foundation-FAPESP and Takeda, as well as the Innovative Medicines Initiative Joint Undertaking ULTRA-DD grant 115766 and the Wellcome Trust (106169/Z/14/Z). L.C. is supported by the Wellcome Trust (OXION grant No: WT084655MA). N.D. is supported by Deutsche Forschungsgemeinschaft (DFG) grant DE1482-4/1.

We thank Diamond Light Source Ltd for access to the macromolecular crystallography beamlines and we thank the Diamond Light Source staff for their help with data collection. We thank all members of the SGC Biotech team, including Shayla Venkaya, Claire Strain-Damerell; Kasia Kupinska, Dong Wang and Katie Ellis. We thank all members of the SGC IMP1 group, including Yin Yao Dong. We thank David Eberhardt for help with the electrophysiology experiments. We are grateful to Rod Chalk, Tiago Moreira, Georgina Berridge and Octawia Borkowska for help with mass spectrometry and Brian Marsden and David Damerell, James Bray, James Crowe and Chris Sluman for bioinformatics support. We thank Frank von Delft, Tobias Krojer and Beth MacLean for assistance with crystallography infrastructure. We thank Oxana Nowak for technical assistance. We thank Thomas Baukrowitz (Christian Albrechts University of Kiel, Kiel, Germany) for critical reading of the manuscript.

## Author Contributions

The project was conceived and managed by E.P.C., N.D. and T.M.. K.E.J.R., obtained crystals that diffracted to beyond 4 Å resolution, collected data and solved the structure of TASK-1 and the complexes with drug molecules, assisted by D.S., and supervised by A.C.W.P. and E.P.C.. W.Z. was involved in optimization of the constructs, protein purification and production of initial crystals, supervised by A.Q. and E.P.C.. A.K.K. performed voltage clamp recordings, pharmacological experiments and single channel recordings and S.R. performed voltage clamp recordings, introduced all TASK-1 mutants and performed ELISA surface expression assays, M.G. performed all experiments with sevoflurane, all experiments were supervised by N.D.. Initial constructs for structural biology were screened for expression by L.S. and large scale insect cell expressions were produced by S.M.M.M., supervised by N.B.B.. Initial tests of the relative activities of TASK-1 and the crystal construct were performed by L.J.C., supervised by S.J.T.. The TASK-1 inhibitors were designed and produced by M.D., M.H., H.M and T.M., supervised by T.M.. Data were analyzed and the paper was written by K.E.J.R., A.C.W.P., A.K.K., S.R., L.C., S.J.T., T.M., N.D. and E.P.C..

## Author Information

M.D. M.H., H.M., T.M., are inventors on patent no. WO2017097792A1, Priority date 10 December, 2018, entitled “2-phenyl-3-(piperazinomethyl)imidazo[1,2-a]pyridine derivatives as blockers of TASK-1 and TASK-3 channels, for the treatment of sleep-related breathing disorders”. The other authors declare no competing interests. They were not involved in the development of the compounds described in this patent and will not benefit from the patent.

---

**Extended Data Fig. 1.**
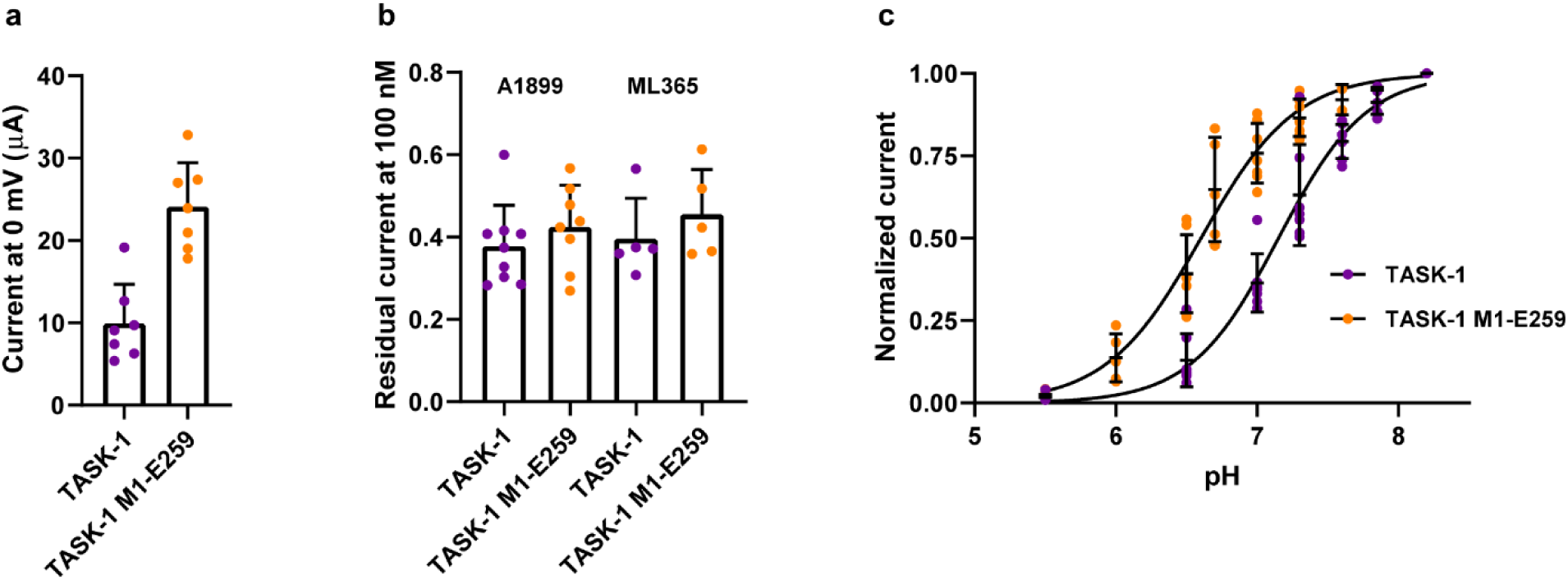
The crystallisation construct TASK-1^M1-E259^ has similar channel properties to WT protein. **a**, Whole-cell TASK-1 wild-type and TASK-1^M1-E259^ currents recorded after expression in *Xenopus* oocytes showing that the crystallisation construct is functionally active. **b**, The truncated crystallisation construct TASK-1^M1-E259^ retains normal inhibition by the pore-blockers A1899 and ML365. **c**, The crystallisation construct retains sensitivity to changes in extracellular pH. The small change in half maximal activation from pH 7.2 to pH 6.4 suggests the truncated cytoplasmic region may influence this process. But overall, the truncated construct exhibits a similar pharmacology and regulation to the wild-type channel. Data are presented as mean ± s.e.m. and shown as individual points.

**Extended Data Fig. 2.**
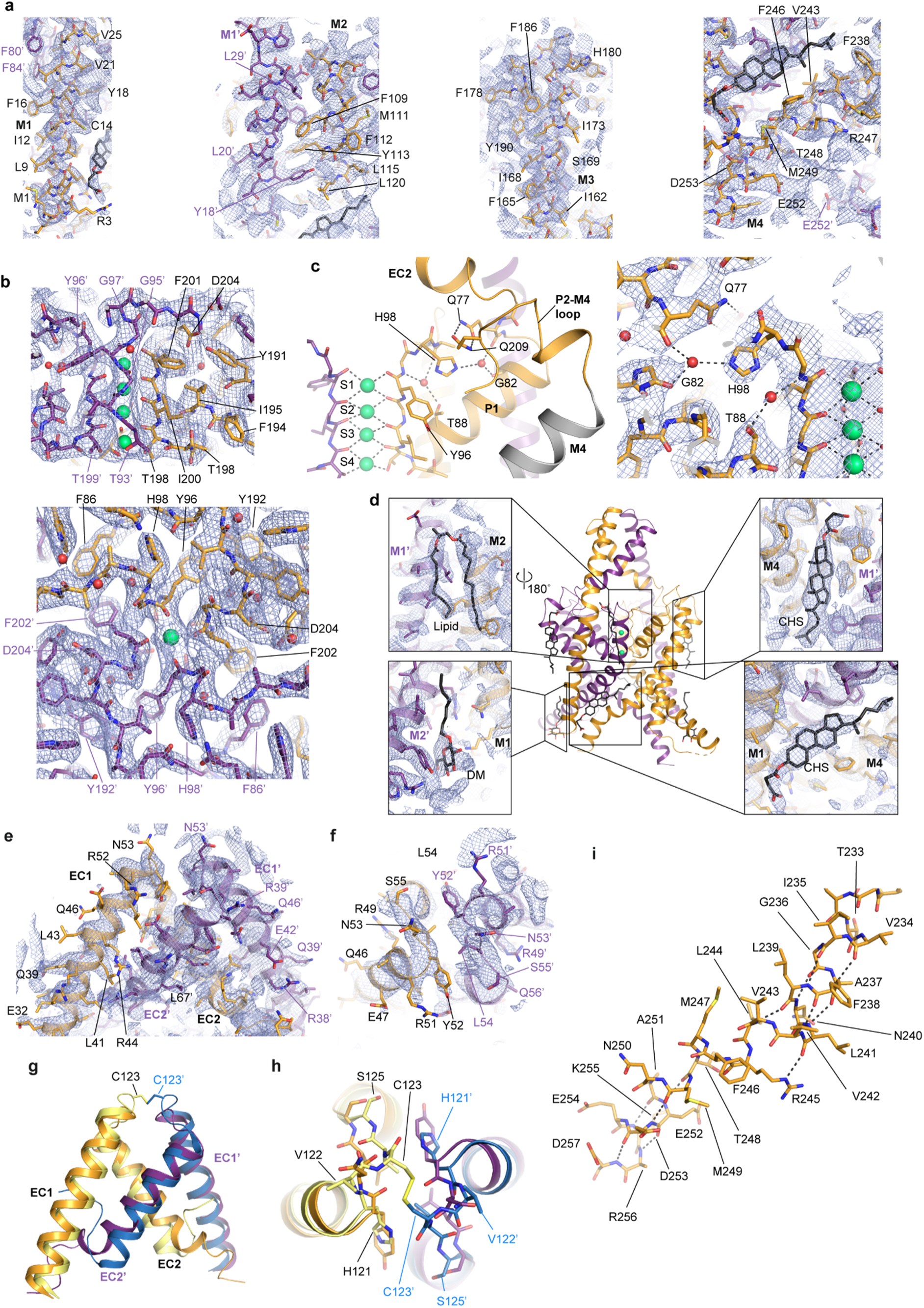
The structure of TASK-1 at 3.1 Å resolution indicates that the cap in TASK-1 forms the classic cap domain structure without an intramolecular disulphide bond. **a**, Final BUSTER 2Fo-Fc electron density maps contoured at 1.0σ, showing the M1-M4 helices. **b**, The selectivity filter with 2Fo-Fc maps shown from the plane of the membrane and from above the membrane. **c**, The pH sensor residue, His98, the surrounding residues and the pore, with the K^+^ ions shown in green, shown both without the map (left) and with the 2Fo-Fc map shown in blue (right). **d**, Cartoon representation of the TASK-1 structure, with lipids, detergents and CHS shown as ball and stick, with zoomed views of the binding sites. **e**, 2Fo-Fc electron density maps of the TASK-1 cap and **f**, view of the cap apex. **g**, superposition of the TASK-1 cap (gold and purple) and TREK-2 (PDB: 4XDJ) cap (yellow and blue), illustrating lack of disulphide bond in TASK-1 and **h**, view of the aligned cap apices. **i**, The M4 helix residues 233-257 represented as sticks, with the main chain hydrogen bonding pattern shown as dashed lines.

**Extended Data Fig. 3.**
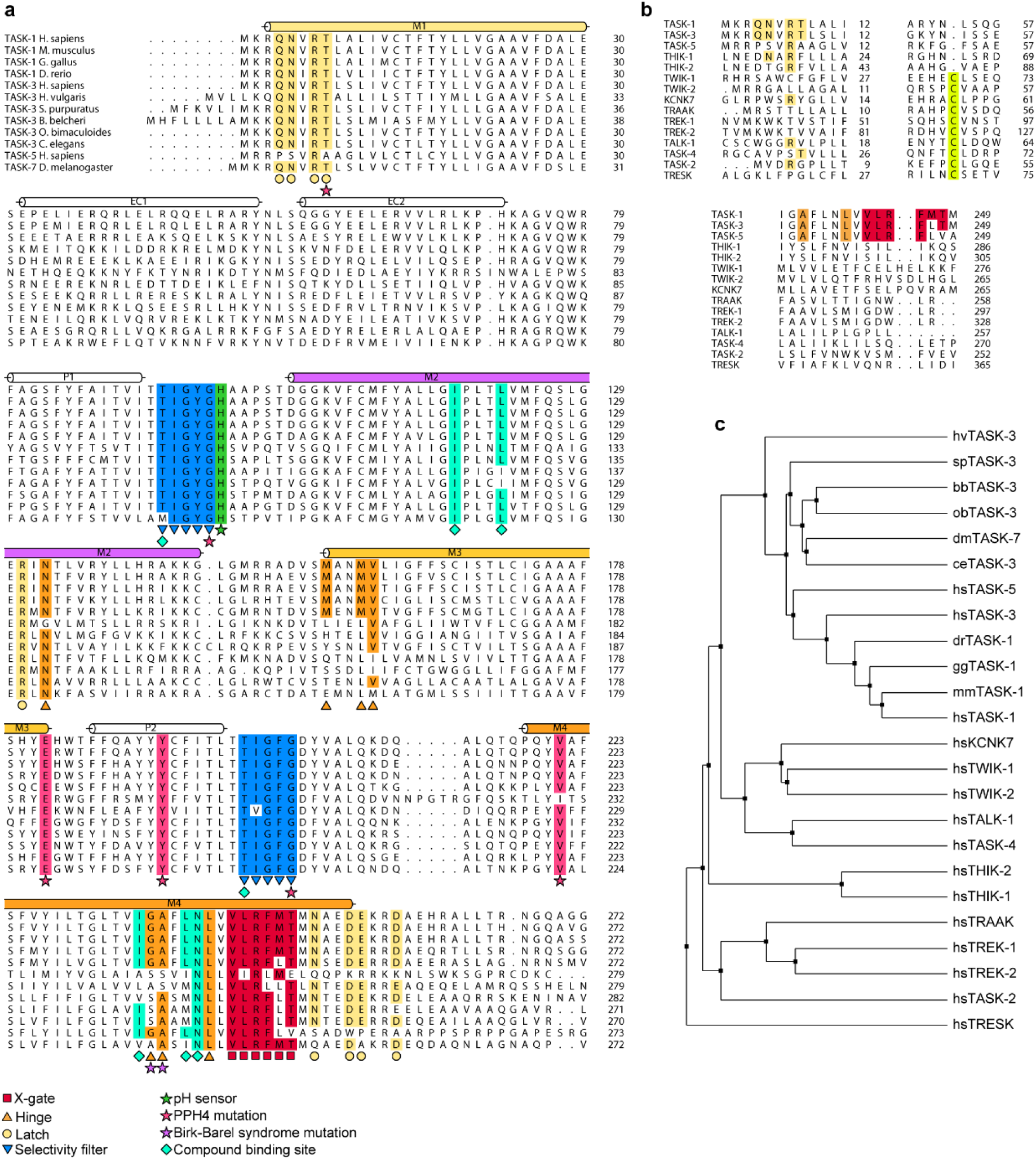
Alignment and phylogenetic trees for the TASKs and K_2P_ channels reveal conservation of X-gate residues in TASK-1, 3 and 5, and a lack of an X-gate in other K_2P_ channels. **a**, Sequence alignment of TASK channels from different species. **b**, Alignment of the human K_2P_ proximal M1, cap and distal M4 regions. The cysteine residue in the cap apex is highlighted in yellow and the X-gate residues in red. **c**, Phylogenetic tree of the human K_2P_s and the TASK channels in **a**. The TASK-1, −3 and −5 channels are highly conserved, whereas the K_2P_ channels that were originally designated at TASK-2 and TASK-4/TALK-2 have much lower homology and belong in the subfamily of TALK channels.

**Extended Data Fig. 4.**
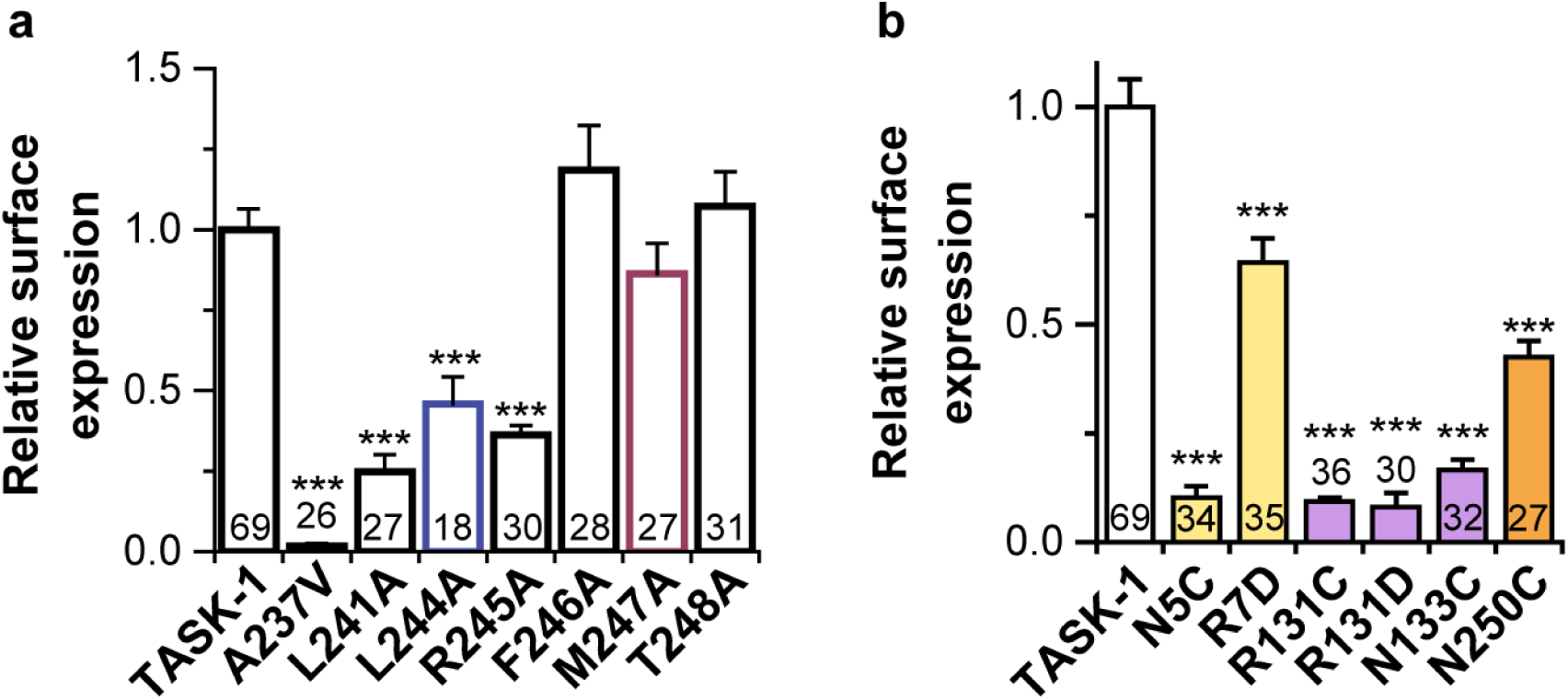
Relative surface expression levels for TASK-1 mutants. **a**, Relative surface expression of wild-type TASK-1 and TASK-1 channels with mutations in the X-gate. **b**, Relative surface expression of wild-type TASK-1 and TASK-1 mutations in the latch region. All data were recorded in *X. laevis* oocytes. Numbers on plots indicate the number of repeats per data point. Data are presented as mean ± s.e.m..

**Extended Data Fig. 5.**
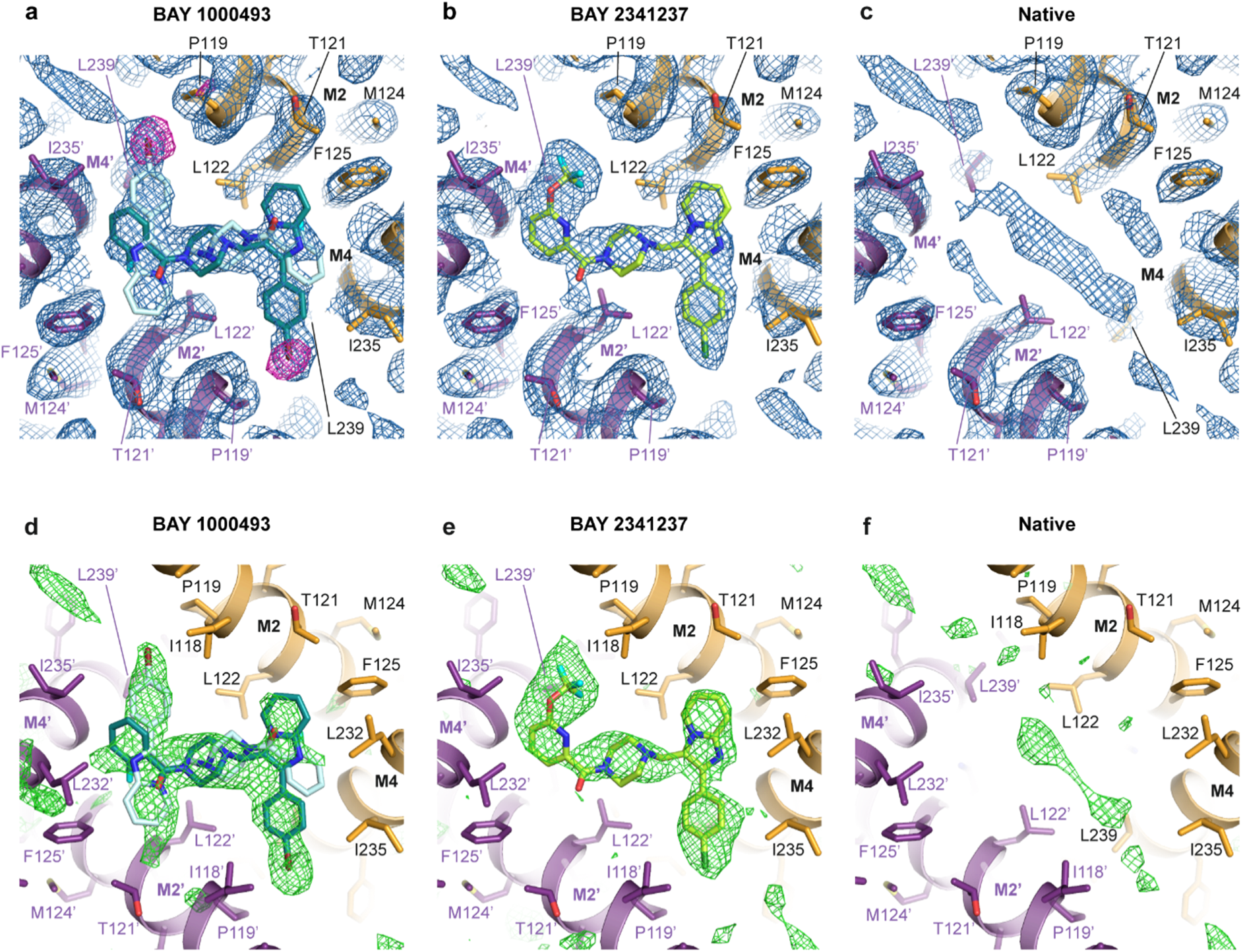
Structures, electron density maps and bromine anomalous data for TASK-1 in complex with BAY 1000493 or BAY 2341237. **a**, The TASK-1/BAY 1000493 complex structure, shown from the selectivity filter, looking towards the vestibule, with the 2Fo-Fc map shown in blue and the anomalous difference map at 3.3 Å collected at the Br edge shown in magenta. The two 50% occupancy BAY 1000493 molecules are shown with carbons coloured teal or light blue, oxygens in red, nitrogens in dark blue, fluorines in cyan and bromines in maroon. **b**, The TASK-1/BAY 2341237 complex structure, shown as for a, with the 2Fo-Fc map shown in blue. The single 100% occupancy BAY 1000493 molecule is shown with carbons coloured green, oxygens in red, nitrogens in dark blue, fluorines in cyan and chlorines in dark green. **c**, The TASK-1 native structure, shown as for a, with the 2Fo-Fc map shown in blue. There is some residual density below the selectivity filter, as is seen in many K2P structures. This type of density is sometimes interpreted as lipid aliphatic chains, but in this instance the correct interpretation was unclear, so we did not model anything into this density. **d, e**, and **f**, represent the same structures as in **a, b** and **c**, with the positive Fo-Fc difference density from omit maps calculated with the inhibitors excluded shown in green. 2Fo-Fc and Fo-Fc maps are contoured at 1.0σ and 2.5σ respectively. The anomalous difference map in **a**, is contoured at 3.5σ.

**Extended Data Fig. 6.**
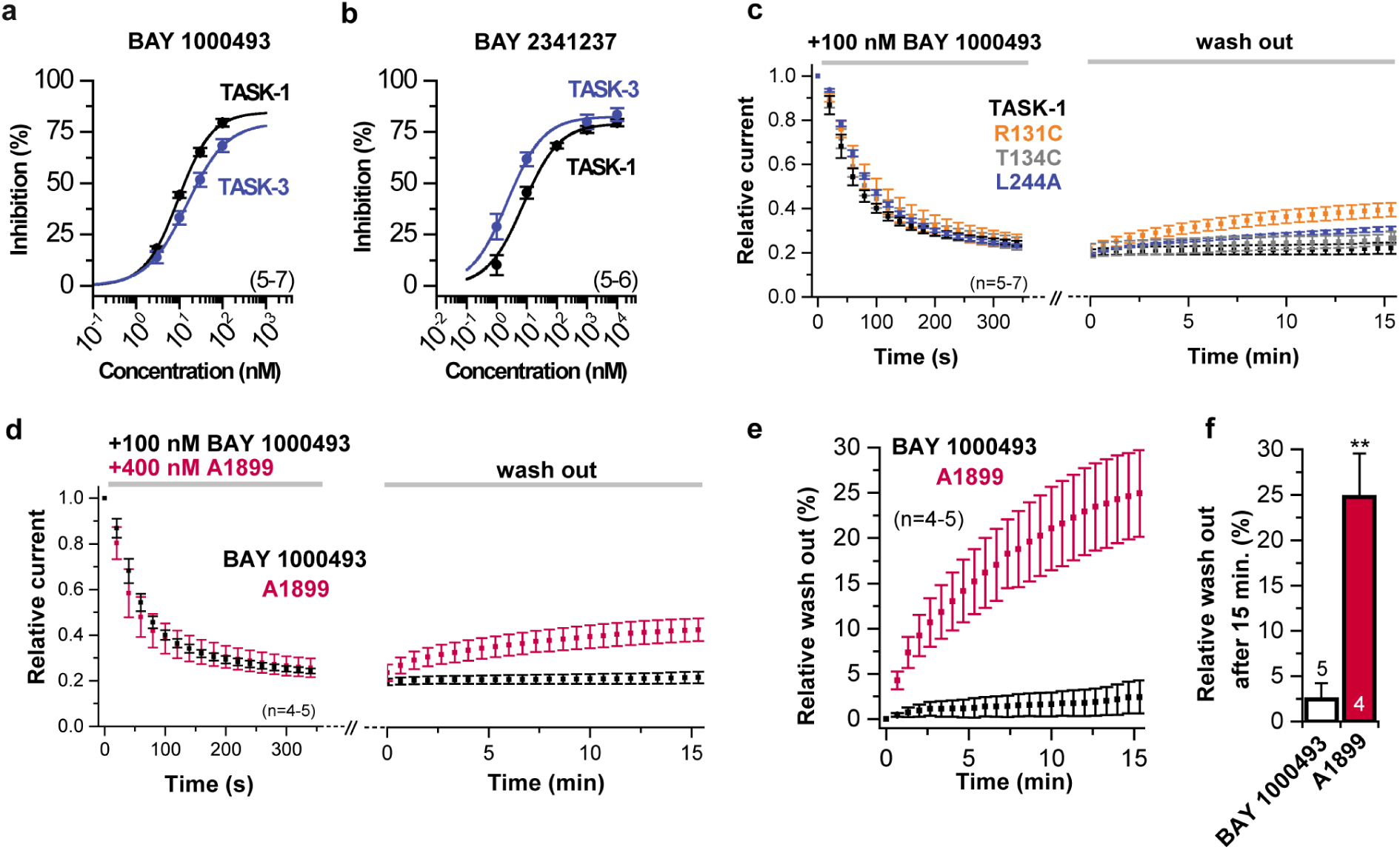
Effects of BAY 1000493 and BAT 2341237 on TASK-1 and TASK-3, and inhibitor washout data for BAY 1000493 and A1899 inhibitor. **a**, Dose response curves for BAY 1000493 on TASK-1 and TASK-3. **b**, Dose response curves for BAY 2341237 on TASK-1 and TASK-3. **c**, Washout of BAY 1000493 from TASK-1 and TASK-1 mutants. **d**, Washout of BAY 1000493 compared to A1899. **e**, relative washout rates for BAY 1000493 and A1899. **f**, Analysis of the washout rates for the compounds shown in **c**. Data are presented as mean ± s.e.m..

**Extended Data Fig. 7.**
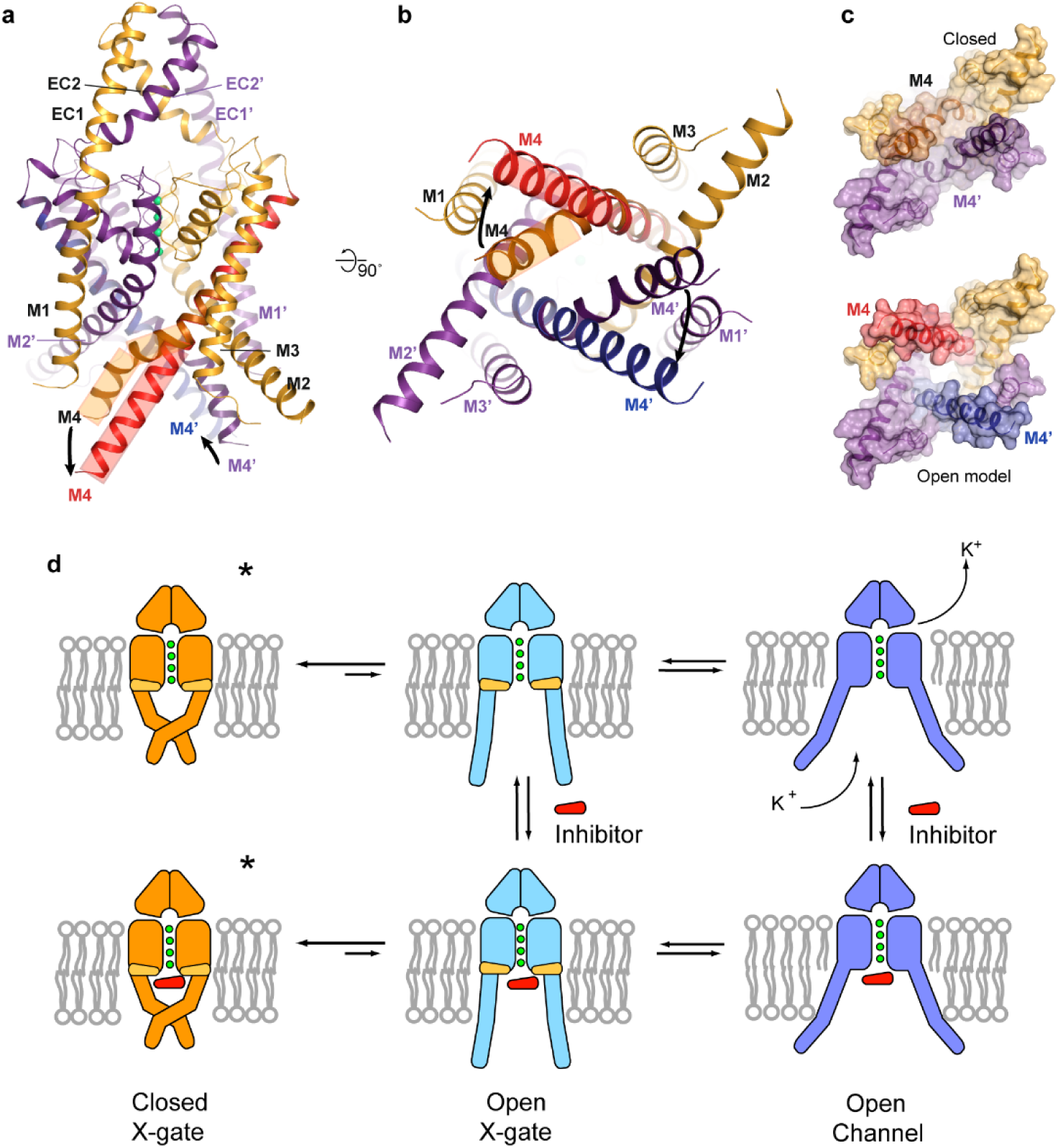
Model of the conformation that TASK-1 could adopt with a straight, continuous M4 helix and a gating scheme for TASK-1. **a,b** Model of TASK-1 with M4 adopting a straight α-helical conformation (in red and blue) aligned with the TASK-1 structure shown from **a**, the side and **b**, below the membrane. **c**, The TASK-1 structure and open model viewed from below the membrane. **d**, A schematic model for TASK-1 activation, with the X-gate closed state shown in orange, the X-gate open state shown in blue and the open channel state shown in mauve. Stars indicate the conformations for which we have structures. The exact nature of the open X-gate and open channel conformations are not yet known.

**Extended Data Fig. 8.**
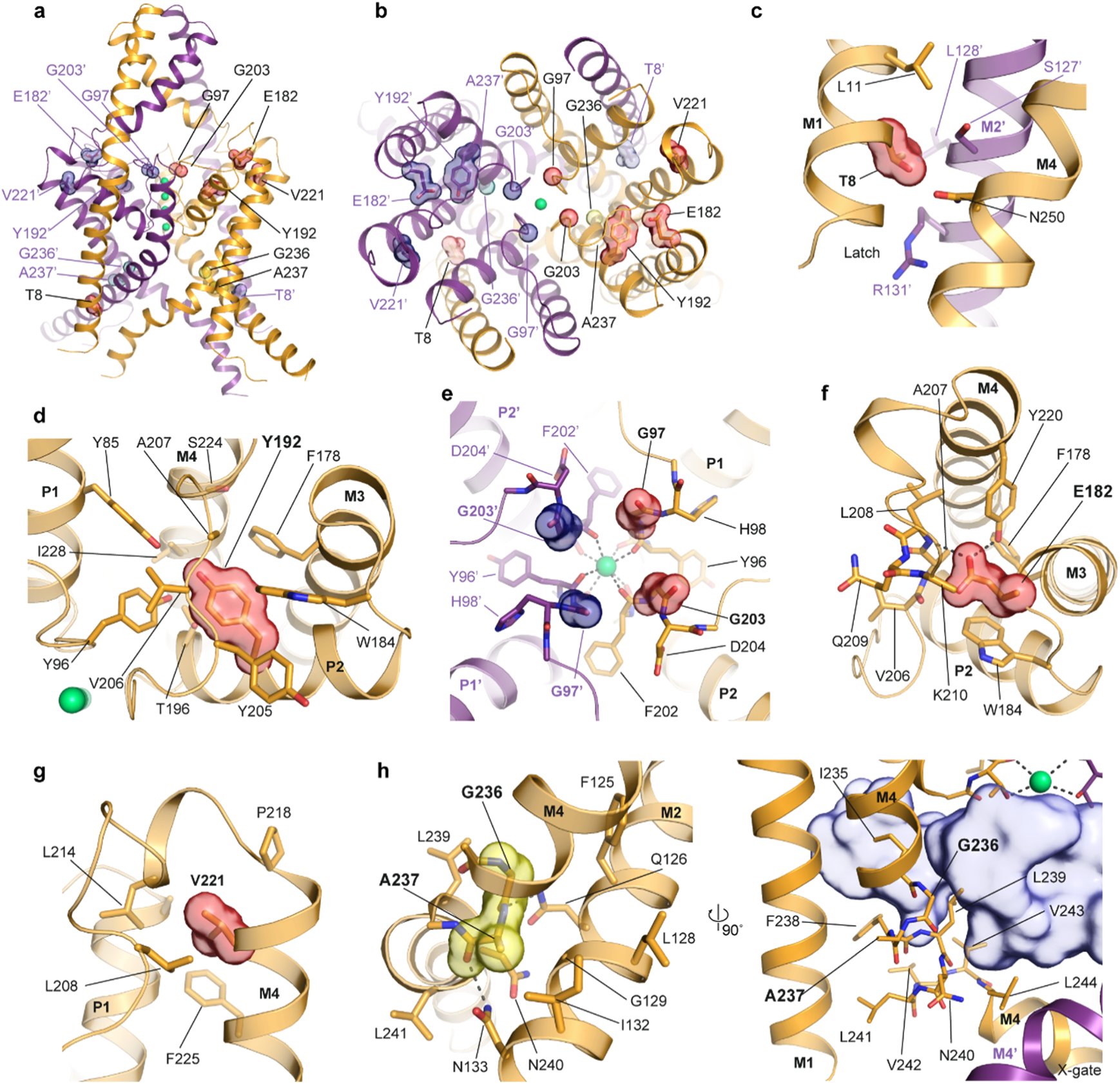
TASK-1 mutations associated with PPH4 – structure and location of mutations. Location of mutations in TASK-1 and TASK-3 mapped onto the TASK-1 structure shown from **a**, the membrane plane and **b**, from the extracellular side. Close-up views of **c** Thr8Lys in the latch region on M1, **d**, Tyr192Cys, **e**, Gly97Arg and Gly203Asp in the pore loops, **f**, Glu182Lys, and **g**,Val221Leu. **h**, the TASK-3 Birk-Barel syndrome mutations Gly236Arg and Ala237Asp in M4 (highlighted in yellow in the panel to the left), shown from the membrane plane and rotated 90° with the vestibule shown in blue in the panel to the right.

**Extended Data Table 1.**
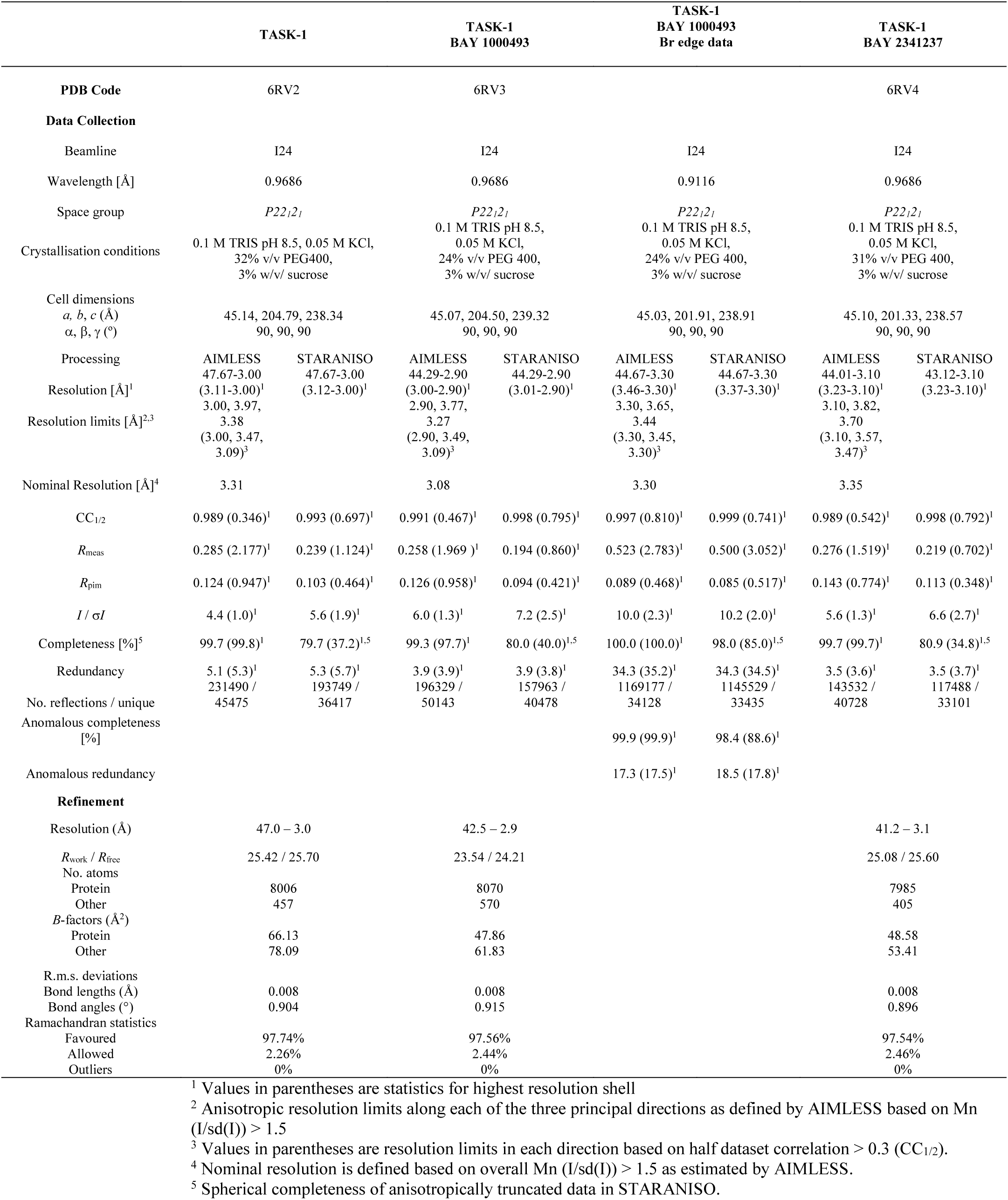
X-ray crystallography data, refinement and model statistics.

**Extended Data Table 2.** Effects of BAY1000493 on a range of ion channels.

| **Voltage sensitive dye measurements**<br>**(FLIPR membrane potential assay)<sup>1</sup>** |  |
| --- | --- |
| TRESK | >30 uM |
| TWIK2 | >30 uM |
| EAG1 | >30 uM |
| K<sub>ir</sub>3.1/3.4 | >30 µM |
| K<sub>v</sub>7.1 | >10 µM |
| K<sub>ca</sub>3.1 | >30 µM |
| **Calcium sensitive dye measurements**<br>**(FLIPR Tetra System)<sup>2</sup>** |  |
| Ca<sub>v</sub>3.2 | >30 µM |
| TRPC5 | >30 µM |
| TRPC6 | >30 µM |
| TRPA1 | >30 µM |
| TRPV4 | >30 µM |
| **Extended Cardiac Safety Profiler Panel (Eurofins)<sup>3</sup>** |  |
| hERG | 2.4 uM |
| Na<sub>v</sub>1.5 | >30 µM |
| Ca<sub>v</sub>1.2 | >30 µM |
| K<sub>v</sub>1.1 | 29 µM |
| K<sub>v</sub>1.3 | 17 µM |
| K<sub>v</sub>1.5 | 19 µM |
| K<sub>v</sub>4.3 | >10 µM |
| KCNQ1 | >30 µM |
| HCN4 | >30 µM |

<sup>1</sup>.As described here: https://www.moleculardevices.com/sites/default/files/en/assets/data-sheets/reagents/flipr-membrane-potential-assay-kits.pdf
<sup>2</sup>.As described here: https://www.moleculardevices.com/sites/default/files/en/assets/data-sheets/reagents/calcium-assays-for-the-flipr-tetra-system.pdf
<sup>3</sup>.As described here: https://www.eurofinsdiscoveryservices.com/cms/cms-content/services/in-vitro-assays/ion-channels/cardiacprofiler/

